# A resource for improved predictions of *Trypanosoma* and *Leishmania* protein three-dimensional structure

**DOI:** 10.1101/2021.09.02.458674

**Authors:** Richard John Wheeler

**Affiliations:** Peter Medawar Building for Pathogen Research, University of Oxford, South Parks Road, Oxford, OX1 3SY. UK

## Abstract

AlphaFold2 and RoseTTAfold represent a transformative advance for predicting protein structure. They are able to make very high-quality predictions given a high-quality alignment of the protein sequence with related proteins. These predictions are now readily available *via* the AlphaFold database of predicted structures and AlphaFold/RoseTTAfold Colaboratory notebooks for custom predictions. However, predictions for some species tend to be lower confidence than model organisms. This includes *Trypanosoma cruzi* and *Leishmania infantum*: important unicellular eukaryotic human parasites in an early-branching eukaryotic lineage. The cause appears to be due to poor sampling of this branch of life in the protein sequences databases used for the AlphaFold database and ColabFold. Here, by comprehensively gathering openly available protein sequence data for species from this lineage, significant improvements to AlphaFold2 protein structure prediction over the AlphaFold database and ColabFold are demonstrated. This is made available as an easy-to-use tool for the parasitology community in the form of Colaboratory notebooks for generating multiple sequence alignments and AlphaFold2 predictions of protein structure for *Trypanosoma*, *Leishmania* and related species.

## Introduction

Machine learning approaches to protein structure prediction have crossed a critical success threshold. While predicting the three-dimensional structure of a protein from sequence alone is still unsolved problem, a multiple sequence alignment (MSA) of the target protein sequence with related proteins provides key additional information. Cutting edge approaches using such MSAs now have the potential to reach very high accuracy. This is the input for AlphaFold2^1^ and RoseTTAfold^2^, with AlphaFold2 reaching the highest accuracy prediction at the most recent CASP competition (CASP14^3^) – an accuracy comparable to experimental protein structure determination. AlphaFold2-predicted structures for the near-whole proteome of 21 species^4^ has been made publicly available, and tools like ColabFold^5^ and the official AlphaFold Colaboratory notebook^6^ make custom predictions easily accessible.

Trypanosomatids pose a challenge because of their large evolutionary distance from common model eukaryotes. This order includes important unicellular human, animal and plant parasites, including the human infective *Trypanosoma cruzi*, *Trypanosoma brucei* and many human-infective *Leishmania* species. *T. cruzi* and *Leishmania infantum* are the most deadly of these species and were included in the initial 21 AlphaFold whole proteome predictions^1,4^. Trypanosomatids are members of an early-branching eukaryote linage (Discoba) which also includes the less common, but still deadly, pathogen *Naegleria fowleri*. Other speciose Discoba lineages are Euglena and Diplonema, unicellular aquatic organisms and important and abundant auto- and heterotrophic plankton respectively. An initial inspection of the AlphaFold database (alphafold.ebi.ac.uk) suggested protein structure prediction accuracy for *T. cruzi* and *L. infantum* is often low – particularly for kinetoplastid specific proteins. Many of these are vital, like the unconventional kinetochore proteins^7^.

Discoba diversity is less well sampled by genomes and transcriptomes than lineages like plants or metazoa, making construction of deep MSAs more difficult. This is important as MSAs encode additional structural information beyond the primary protein sequence alone: They capture evidence for co-evolution of different regions of the primary sequence which may correspond to proximity or interaction in the three-dimensional structure. AlphaFold2/RoseTTAFold prediction of protein structure is greatly improved by high MSA quality and depth, with high MSA coverage critical for high confidence predictions^1,2^. While new approaches^8^ are trying to move beyond multiple sequence alignments, MSAs will remain a powerful source of information.

Currently, the input databases for the AlphaFold database and the ColabFold notebook are of UniRef^9^ plus environmental sample sequence databases (BFD, Uniclust and MGNify^10–12^). However, these have relatively poor coverage of Discoba. It appears that a significant quantity of genomic and transcriptomic data available in the community genome resource TriTrypDB^13,14^, the NCBI genome^15^, transcriptome shotgun assembly (TSA)^16^ and sequencing read archive (SRA)^17^ databases are not incorporated. It seemed likely that this is simple opportunity to improve protein MSAs for protein structure predictions for *Trypanosoma*, *Leishmania* and other Discoba species.

Here, this additional protein sequence data was gathered into a comprehensive Discoba database and the ColabFold MMSeqs2-based pipeline^5,18^ was modified to also include the result of a HMMER search of Discoba. Using a test set of 30 *L. infantum* proteins, MSA coverage was always improved, leading to increased AlphaFold2 prediction accuracy in 2/3 of cases. Improvements were greatest for kinetoplastids-specific proteins, with dramatic improvements often possible. The necessary tools to make similar protein structure predictions have been made openly available: The Discoba protein sequence database (for custom searches and MSA generation), Colaboratory notebooks for generating MSAs by HMMER or MMSeqs2 (for use in AlphaFold2 or RoseTTAFold implementations), and a standalone Colaboratory notebook for AlphaFold2 structure predictions based on ColabFold incorporating a search of this additional database. These are available at github.com/zephyris/discoba_alphafold.

## Methods

Predicted protein sequences were gathered from 248 Discoba transcriptomes or genomes (Table S1): 162 genomes and 86 transcriptomes. 157 from cultured populations (almost all axenic) and 91 from single cell samples. Currently, 208 are included in the gathered Discoba protein sequence database, with the remainder pending assembly optimisation.

### Genome-derived

TriTrypDB: All 53 trypanosomatid species available in TriTrypDB^13,14^ release 53, using the provided predicted protein sequences on TriTrypDB where available. For the 17 without predicted protein sequences the translation of all predicted open reading frames (ORFs) over 100 amino acids were used, as kinetoplastids have compact genomes with short intergenic sequences and extremely low occurrence of introns.

NCBI Genomes: 32 genomes for Discoba species. For the 14 with predicted protein sequences on NCBI the existing prediction was used. For the 18 without predictions, the translation of all ORFs over 100 amino acids were used.

Sequencing read archive (SRA): 19 whole genome sequencing (WG-seq) datasets from axenic cultures of Discoba species and 32 single cell WG-seq datasets. For each, assembly was carried out using Velvet^19,20^ (see Genome assembly) and all predicted ORFs over 100 amino acids were used.

### RNA-seq derived

Transcriptome shotgun assembly (TSA) database: 12 transcriptomes for Discoba species, using protein sequence predicted by TransDecoder^21^.

Marine Microbial Eukaryotic Transcriptome Sequencing Project (MMETSP): 2 transcriptomes for Discoba species, using the provided protein sequences which were predicted using TransDecoder.

NCBI SRA: 19 mRNA-seq datasets from axenic cultures of Discoba species, 3 mRNA-seq datasets from mixed cultures including a Discoba species and 59 single cell mRNA-seq datasets. For each, transcriptome assembly was carried out using Trinity^22–24^ followed by protein sequence prediction with TransDecoder^21^ (see Transcriptome assembly).

### Transcriptome assembly

Transcriptome assembly from RNA-seq data used a standardised pipeline, with the same approach used for axenic culture, mixed culture and single cell transcriptomic data. Reads were first error corrected using Rcorrector v1.0.4^25,26^ (using Jellyfish v2.3.0^27,28^) and corrected reads tidied using TranscriptomeAssemblyTools^29^. Any remaining adaptor sequences were trimmed using TrimGalore v0.6.0^30,31^ (using Cutadapt v2.8^32,33^) then an assembly was generated using Trinity v2.12.0^22–24^. Many of these species use polycistronic transcription with a single spliced leader sequence trans-spliced onto the start of all mRNAs. As such common sequences may affect assembly, a two-step approach was used. First, a trial assembly using 1,000,000 reads (or all reads if fewer were available) was generated and the common spliced leader sequence identified using a custom script. Cutadapt was then used to trim reads to remove the spliced leader, then a final assembly was generated using 40,000,000 reads (or all reads if fewer were available). Very similar transcript sequences were removed using cd-hit-est v4.8.1 (part of CD-HIT^34,35^) then translated to predicted proteins using TransDecoder v5.5.0^21^ LongOrfs.

### Genome assembly

Genome assembly from WG-seq data also used a standardised pipeline. For single cell genomic data, reads were first error corrected using Rcorrector v1.0.4^25,26^ (using Jellyfish v2.3.0^27,28^) and TranscriptomeAssemblyTools^29^. For all assemblies, any remaining adaptor sequences were trimmed using TrimGalore v0.6.0^30,31^ (using Cutadapt v2.8^32,33^) then an assembly was generated using Velvet v1.2.10^19,20^ using all available reads. As expected coverage and insert size are not necessarily known, a refinement step was used. Reads were aligned to the assembly using bwa mem v0.7.17^36,37^ and insert size and mean coverage determined using samtools v1.10^38,39^, then a final assembly was generated using Velvet including these parameters and a minimum coverage threshold of 0.25 the mean trial assembly coverage. All open reading frames ≥300 bp (all three frames, both strands) were identified using a custom script.

### Orthology

Protein orthogroups were identified using OrthoFinder v2.5.4^40–42^ (using diamond v2.0.5.143^43–45^ and FastME 2.1.4^46^). Reciprocal best protein sequence search hits were carried out using diamond v2.0.5.143^43–45^ with no e-value cut-off. OrthoFinder and reciprocal best sequence search hit analysis were carried out on a diverse set of 77 UniProt reference eukaryote proteomes^47^: UP000001450, UP000002729, UP000007800, UP000012073, UP000054560, UP000000437, UP000001542, UP000005203, UP000008144, UP000013827, UP000059680, UP000000539, UP000001548, UP000005226, UP000008153, UP000014760, UP000179807, UP000000559, UP000001593, UP000005640, UP000008493, UP000018208, UP000186817, UP000000560, UP000001926, UP000006548, UP000008524, UP000023152, UP000218209, UP000000561, UP000001940, UP000006671, UP000008743, UP000027080, UP000247409, UP000000589, UP000001950, UP000006727, UP000008827, UP000030693, UP000265515, UP000000600, UP000002195, UP000006729, UP000009022, UP000030746, UP000265618, UP000000759, UP000002296, UP000006906, UP000009138, UP000036983, UP000316726, UP000000803, UP000002311, UP000007110, UP000009168, UP000037460, UP000323011, UP000000819, UP000002485, UP000007241, UP000009170, UP000051952, UP000324585, UP000001357, UP000002494, UP000007305, UP000009377, UP000054408, UP000444721, UP000001449, UP000002640, UP000007799, UP000011083, UP000054558. This includes 6 kinetoplastid species (*Bodo saltans*, *Leishmania infantum*, *Leishmania mexicana*, *Perkinsela sp*., *Trypanosoma brucei brucei* and *Trypanosoma cruzi*), which were used as the basis for identifying kinetoplastid specific proteins.

### Intrinsically disordered domains

Intrinsically disordered domains were predicted using IUPred2A^48^ using a score threshold of 0.5 for classification of a residue as disordered.

### AlphaFold2 predictions

Existing AlphaFold predictions of protein structures for *Leishmania infantum* (UP000008153) *Trypanosoma cruzi* (UP000002296) and *Mus musculus* (UP000000589) were taken from alphafold.ebi.ac.uk^1^, last updated using AlphaFold v2.0 2021-07-01. Per residue and global pLDDT was taken from the mmCIF file, PAE from the error json file.

ColabFold predictions were made using an unmodified version of ColabFold^5,18^, with the default MSA pipeline, a MMseqs2^49,49^ search of UniRef^9^ and environmental sample sequence databases^10–12^. Predictions were done using AlphaFold2 parameters from 2021-07-14, not using Amber^51^ relaxation and not using PDB^52^ templates. Due to GPU memory availability in Goole Colaboratory, predictions were restricted to proteins with ≤800 amino acids.

AlphaFold2 predictions incorporating the new Discoba protein sequence database were carried out using a modified version of ColabFold^5,18^. The MMseqs2^49,49^ search of UniRef^9^ and environmental sample sequence databases^10–12^, was supplemented with a HMMER (part of HH-suite)^50^ search of the Discoba protein database described here. A MMSeqs2 search of the Discoba protein database was also trialled, but use of HMMER for Discoba searches typically gave a small increase in pLDDT, presumably as AlphaFold v2.0 was trained using HMMER MSAs. Predictions were again done using AlphaFold2 parameters from 2021-07-14, not using Amber relaxation and not using PDB templates.

A test set of *L. infantum* proteins were selected randomly, see Results. Randomly selected conserved genes: A4HU53, A4I2E1, A4HUD2, A4IA46, A4I444, A4IC57, A4IAB2, A4I7M6, A4I0C5, A4HTD2. Randomly selected not conserved genes: A4I944, A4HYM2, A4I1S6, A4I5D0, A4IAG0, A4I787, A4I9X8, A4IB72, A4I5C1, A4HS18. Randomly selected not conserved ‘promising’ genes: E9AGZ8, A4HW74, A4HZS9, A4I0P7, A4I2Z9, A4IDS7, A4HRK9, A4IBK2, A4I4D7, E9AGB8.

## Results

Many AlphaFold2-predicted protein structures for *Leishmania infantum* and *Trypanosoma cruzi* report high predicted local distance difference test score (pLDDT)^53^, a per residue 0 to 100 score with high values showing higher confidence, and low predicted average error (PAE), a per residue pair distance score with low values showing lower error. Using the Alphafold database pLDDT^1,4^, proteome-wide performance of AlphaFold2 can be evaluated quantitatively. For comparison mouse (*Mus musculus*) was selected as, unlike human proteins, predictions were carried out without special treatment. Overall, *L. infantum* and *T. cruzi* protein structure predictions are skewed to lower pLDDT (Fig. 1A).

**Fig. 1.**
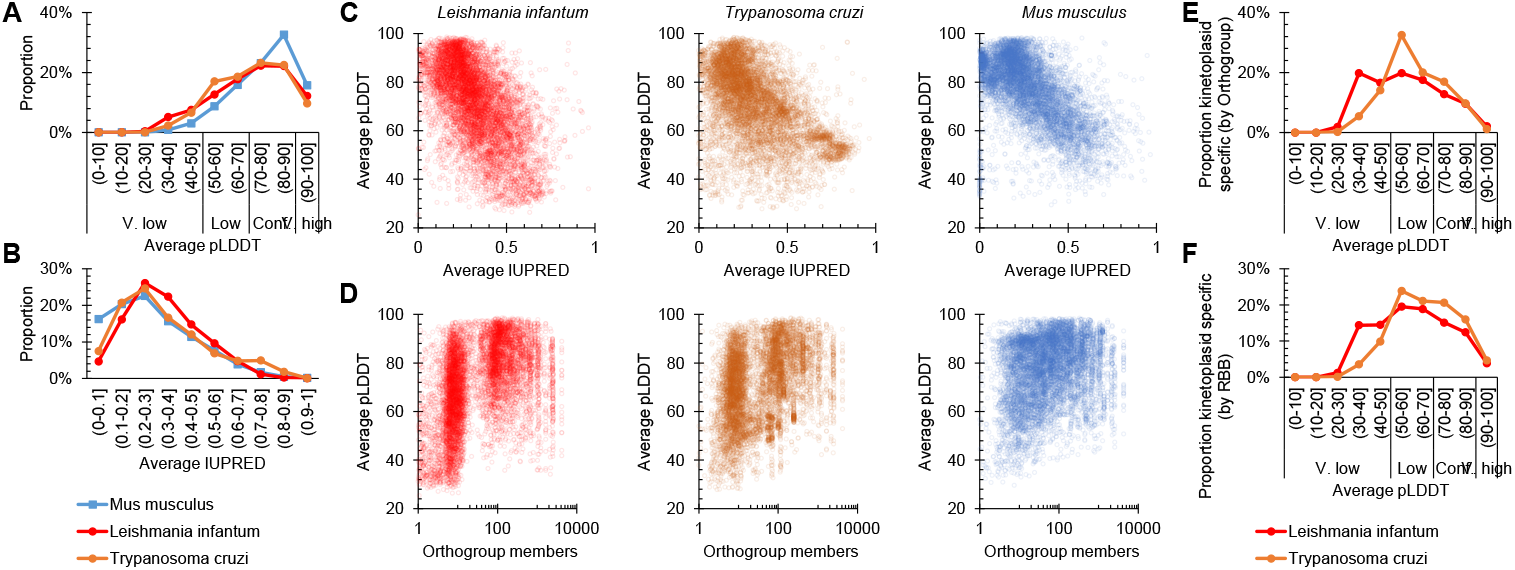
Proteome-wide quality of protein structure predictions of kinetoplastid proteins in comparison to mouse proteins in the AlphaFold database. **A)** Distribution of per-protein average pLDDT for all *L. infantum* (7924), *T. cruzi* (19024) and, for comparison, *M. musculus* (21588) proteins, from the AlphaFold database^1,4^. Scores for very low, low, confident and very high confidence categories are the same as used on the website. **B)** Distribution of per-protein average IUPred score for the same three species. **C)** Correlation of per-protein average pLDDT with IUPred score for the same three species. **D)** Correlation of per-protein average pLDDT with number of orthologs (total number of orthogroup members determined from a diverse set of eukaryotes, see Methods). A random number between 0 and 1 was added to each ortholog count to better represent point density at low ortholog numbers. **E,F)** Distribution of per-protein average pLDDT for all *L. infantum* and *T. cruzi* proteins lacking an ortholog outside of the kinetoplastids, as determined by either **E)** orthogroup members only in kinetoplastid species (1509 and 7181 proteins respectively) or **F)** reciprocal best protein sequence search hits only in kinetoplastids species (2361 and 11723 proteins respectively).

Lower pLDDT could be explained by more disordered protein domains, as predictions for these regions correlate with pLDDT^1^. *L. infantum* and *T. cruzi* proteins do not have a markedly different predicted degree of disorder to *M. musculus* (Fig. 1B) although, as expected^1^, pLDDT had a negative correlation with disorder score in all three species (Fig. 1C). Alternatively, it may be a limitation due to the depth of the input protein MSAs. pLDDT correlated with number of orthologs detected using OrthoFinder^40–42^ on a set of 77 reference proteins of diverse eukaryotes (Fig. 1D), confirming this expectation.

Unlike *M. musculus*, the distribution of number of orthologs for *L. infantum* and *T. cruzi* was strongly bimodal with many having fewer than 10. These proteins had, on average, markedly lower pLDDTs (Fig. 1D). Analysis using more stringent measures of protein specificity/uniqueness to the kinetoplastids showed a similar pattern: kinetoplastids-specific proteins were identified as those with only reciprocal best sequence search hits among the kinetoplastids (Fig. 1E) or those with only orthogroup members among the kinetoplastids (Fig. 1F), and both sets had low pLDDT (Fig. 1E,F). Overall, this confirms that MSA quality is likely the limiting factor for many *T. cruzi* and *L. infantum* protein structure predictions.

As much protein sequence data as possible was therefore gathered for Discoba species, drawing upon both TriTrypDB^13,14^ (well known to the *Trypanosoma/Leishmania* community but not entirely in UniProt), and lesser known, unpublished or very recent data available *via* nucleotide sequencing databases, gathered using the NCBI taxonomy browser (see Methods, Table S1). Unlike many applications, precise knowledge about sample/species identity, high sample purity and high transcriptome/genome coverage are not critical – therefore the gather was as inclusive as possible. So far, this includes 208 predicted proteomes deemed high enough quality, representing 1.4 billion amino acids across 4.0 million protein sequences. This may increase with ongoing optimisation.

To benchmark any improvements over the AlphaFold database predictions a set of 30 *L. infantum* proteins were selected: 10 random proteins which have orthologs in many diverse eukaryotes, 10 random proteins which appear unique to the kinetoplastid lineage (no orthogroup members outside the kinetoplastids) and 10 random proteins which appeared ‘promising’ but with a low pLDDT in the AlphaFold database. The latter were selected based on size (avoiding small proteins, ≳300 amino acids), lack of low complexity or repetitive regions (≲30% unstructured and manually avoiding repeats), orthologs in few species (≲10), without numerous paralogs, and low average pLDDT (≲60).

To carry out AlphaFold2 protein structure predictions ColabFold was selected as a fast but high accuracy and accessible AlphaFold2 implementation^5,18^. As expected, unmodified ColabFold gave per-protein mean pLDDTs comparable to, but on average slightly lower than, the AlphaFold database for the test proteins (Fig. 2A). ColabFold was then modified to generate a HMMER-generated MSA from the Discoba database and append this to the default MSA, before carrying out the AlphaFold2 prediction. This improved mean pLDDT for a large majority of protein structure predictions, whether compared to the AlphaFold database or unmodified ColabFold, with less confident predictions seeing the largest improvement (Fig. 2B,C). pLDDT increase occurred at all confidence levels within a protein. Using the confidence thresholds in the AlphaFold database, the proportion of residues over the threshold for a low confidence (>50), confident (>70) and high confidence (>90) prediction almost all increased for a large majority of proteins (Fig. 2D).

**Fig. 2.**
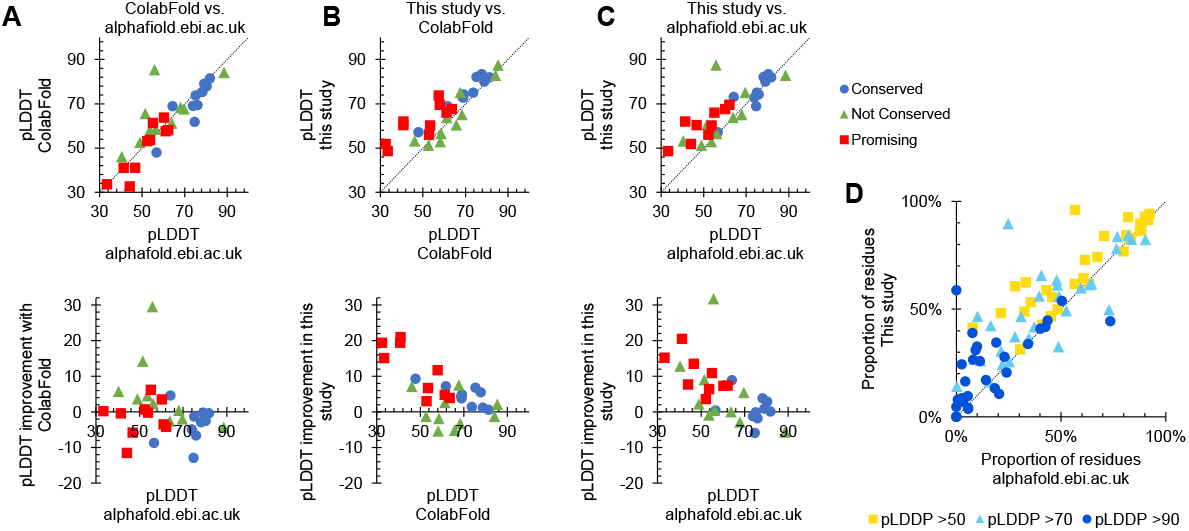
Improved AlphaFold2 predictions using ColabFold and a wider set of Discoba sequences for MSAs. **A-C)** Comparison of per-protein average pLDDT for 30 test proteins, 10 random widely conserved proteins, 10 random kinetoplastids-specific proteins and 10 kinetoplastid-specific proteins which appeared likely to improve with additional MSA sequences. **A)** Unmodified ColabFold in comparison to the AlphaFold database plotted as: Top, raw pLDDTs. Points to the top left of the diagonal represent improved (higher pLDDT) predictions. Bottom, change in pLDDT. Points above the horizontal axis represent improvement. **B)** This study (ColabFold with HMMER search of additional Discoba sequences) in comparison to unmodified ColabFold. **C)** This study in comparison to the AlphaFold database. **D)** The same comparison as C) but plotting the proportion of residues over different threshold pLDDT values instead of mean pLDDT.

Improvement was most marked among the test proteins not conserved outside of the kinetoplastids, especially the ones selected as ‘promising’ (Fig. 2A). Inspection of these showed a range of improvements, including overall large decreases in PAE (Fig. 3A), the first high confidence prediction of any folds (Fig. 3B) and the prediction of a single globular domain rather than two subdomains (Fig. 3C).

**Fig. 3.**
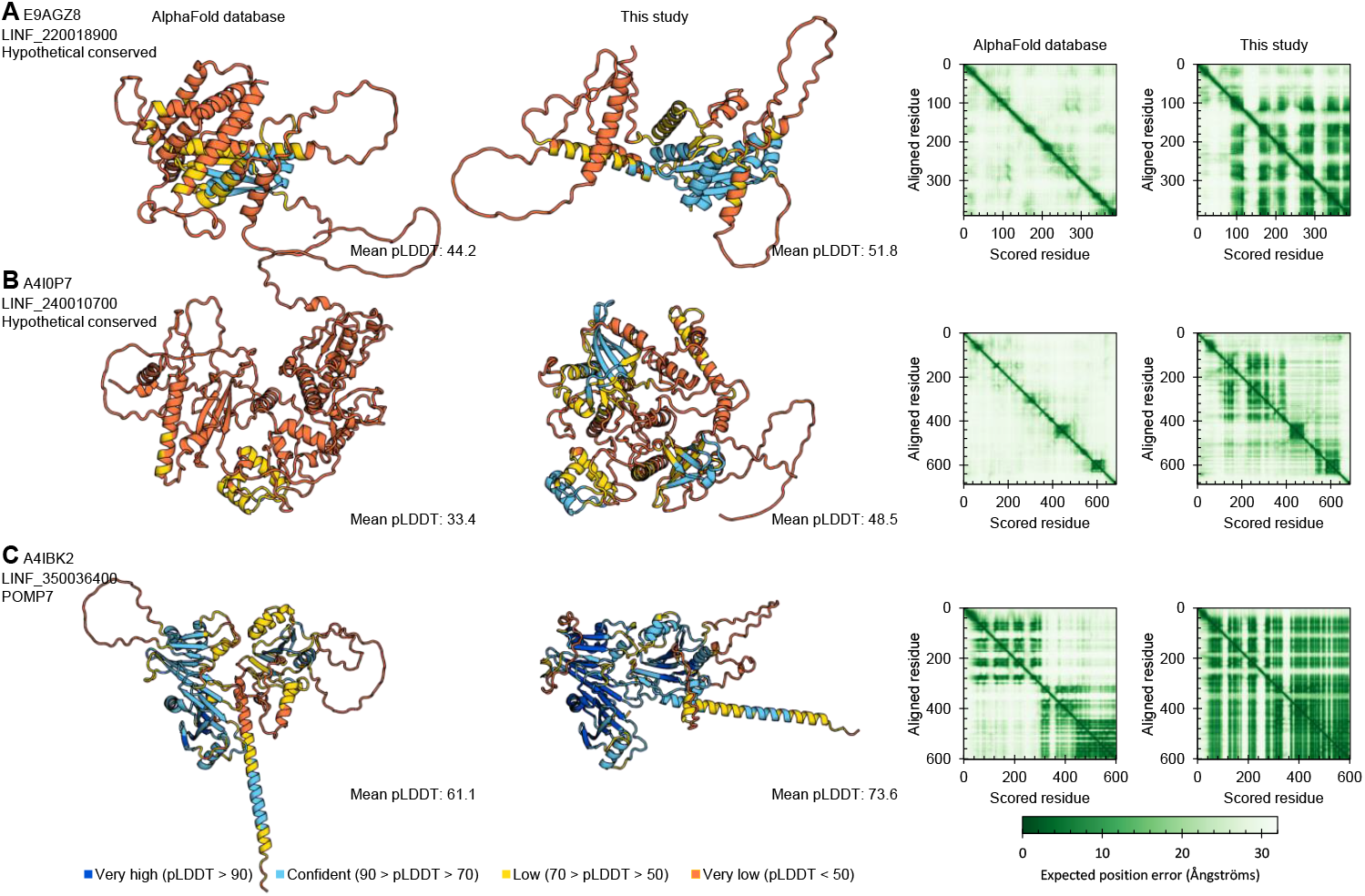
Example *L. infantum* proteins showing significant improvement in structure prediction over the AlphaFold database. Predicted protein structures for three example *L. infantum* proteins showing, from left to right, the AlphaFold database structure, the predicted structure using ColabFold supplemented with a HMMER search of the Discoba database described in this study, the pairwise PAE for the AlphaFold database structure and the PAE for the structure predicted in this study.

## Discussion

This work shows that significant improvement in the pLDDT and PAE of AlphaFold2 structure predictions is possible for *Trypanosoma* and *Leishmania* proteins, relative to the publicly available AlphaFold database at alphafold.ebi.ac.uk^1,4^ and the open tool ColabFold^5,18^ (Fig. 2). This is simply by designing a protein sequence database for MSA generation more appropriate for this branch of eukaryotic life. Easy-to-use tools for MSA generation and AlphaFold2 structure prediction exploiting this Discoba protein sequence database have been made available at github.com/zephyris/discoba_alphafold. Even in these early-branching eukaryotes, the huge advance AlphaFold (and RoseTTAfold) represent can therefore, to a great extent, translate protein structure determination into a genome and transcriptome sequencing problem. Although, experimental protein structure determination will continue to be vital to confirm predictions, explore dynamics and complexes, etc.

Structure prediction improvement was most marked for proteins specific to the kinetoplastids. A large proportion of trypanosomatid parasites’ genomes falls into this group – several hundred to thousands depending on definition (Fig. 1E,F). Many of these proteins lack any domains detectable by primary sequence (sometimes called the ‘dark proteome’) making a structure prediction a first insight into potential function. However, improvement in protein structure prediction at all levels are valuable. It may allow a high-confidence prediction of vital kinetoplastid proteins with orthologs in many species, allowing analysis of high specificity small molecule docking. This is of potential importance for drug development.

Improvement is certainly not guaranteed for any individual protein: Proteins well conserved across diverse eukaryotes will already have deep MSAs giving high confidence structural prediction (*cf*. Fig. 1D). Proteins which have intrinsically disordered domains (eg. many RNA binding proteins) or only gain structure as part of a multisubunit structure (eg. many ribosome proteins) are unlikely to see significant improvement (*cf*. Fig. 1C). Also, proteins which are extremely fast-evolving, or recent innovation found only in a few species, are less likely to benefit.

Properties of kinetoplastids chromosome organisation may enable future developments. The order of genes on chromosomes is well conserved^54^ and, sometimes, this provides additional information which allows identification of orthologs which are difficult or impossible to detect based on primary sequence alone (eg. Basalin^55^). Gathering such syntenic genes allows generation of MSAs of extremely divergent orthologs, although initial tests did not show any benefit for structural prediction – perhaps as AlphaFold2 was not trained on this type of MSA, alternatively the orthologs may simply be too divergent.

Overall, this highlights both the importance of sequencing diverse organisms, for example animal pathogens related to human pathogens and non-pathogenic relatives, and ensuring this data is made available through nucleotide sequencing, genome and proteome databases. It also emphasises that protein sequence data need to be carefully gathered before embarking on important or large-scale structural predictions: Even carefully selected representative databases often retain biases towards model organisms. Similar protein structure prediction improvements are likely possible for other branches of eukaryotic life.

## Acknowledgements

Richard Wheeler is supported by the Wellcome Trust [211075/Z/18/Z]. I would like to particularly thank the numerous research groups who have made their sequencing data available through publicly available databases.

---

**Table S1.**
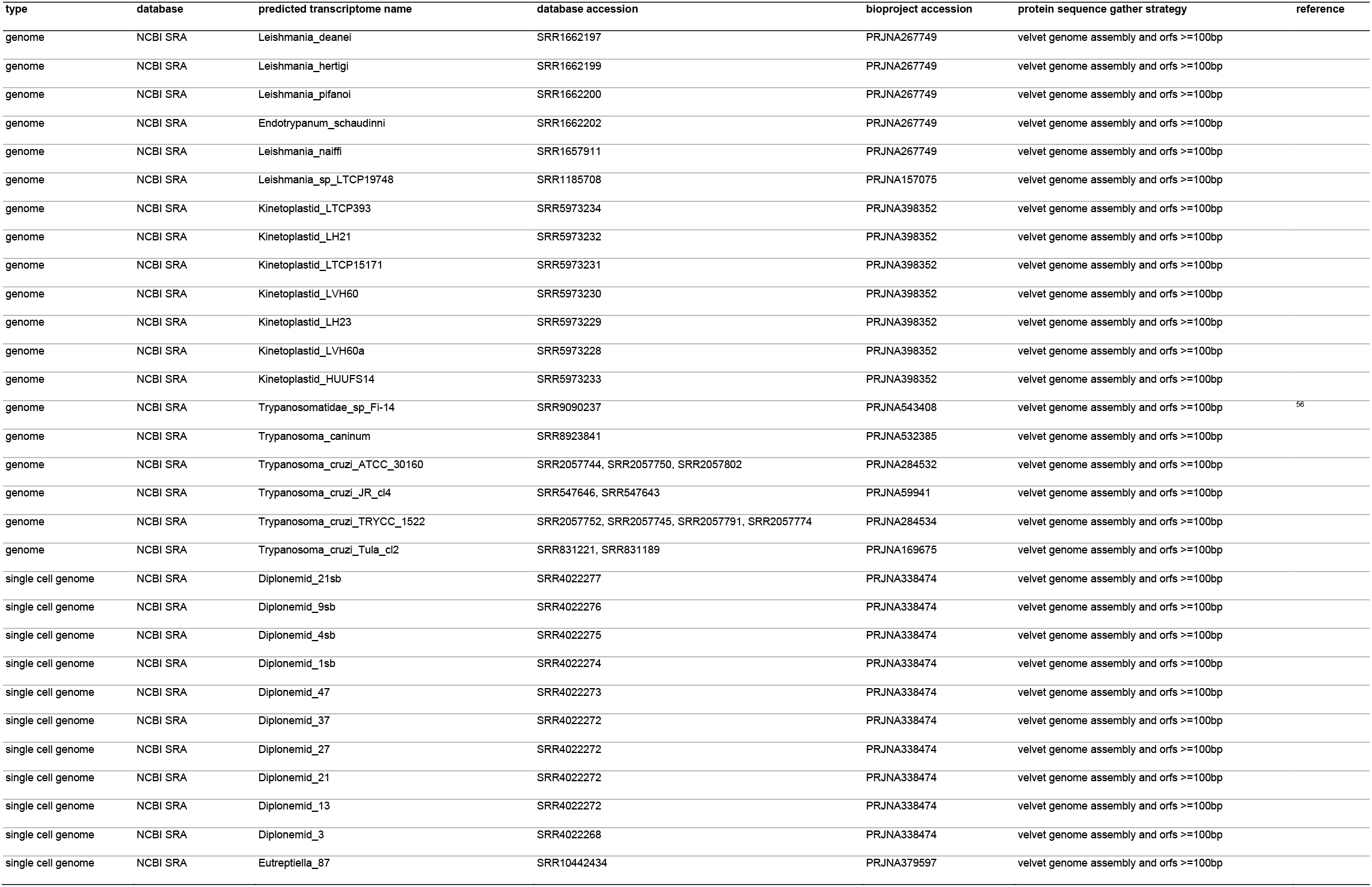
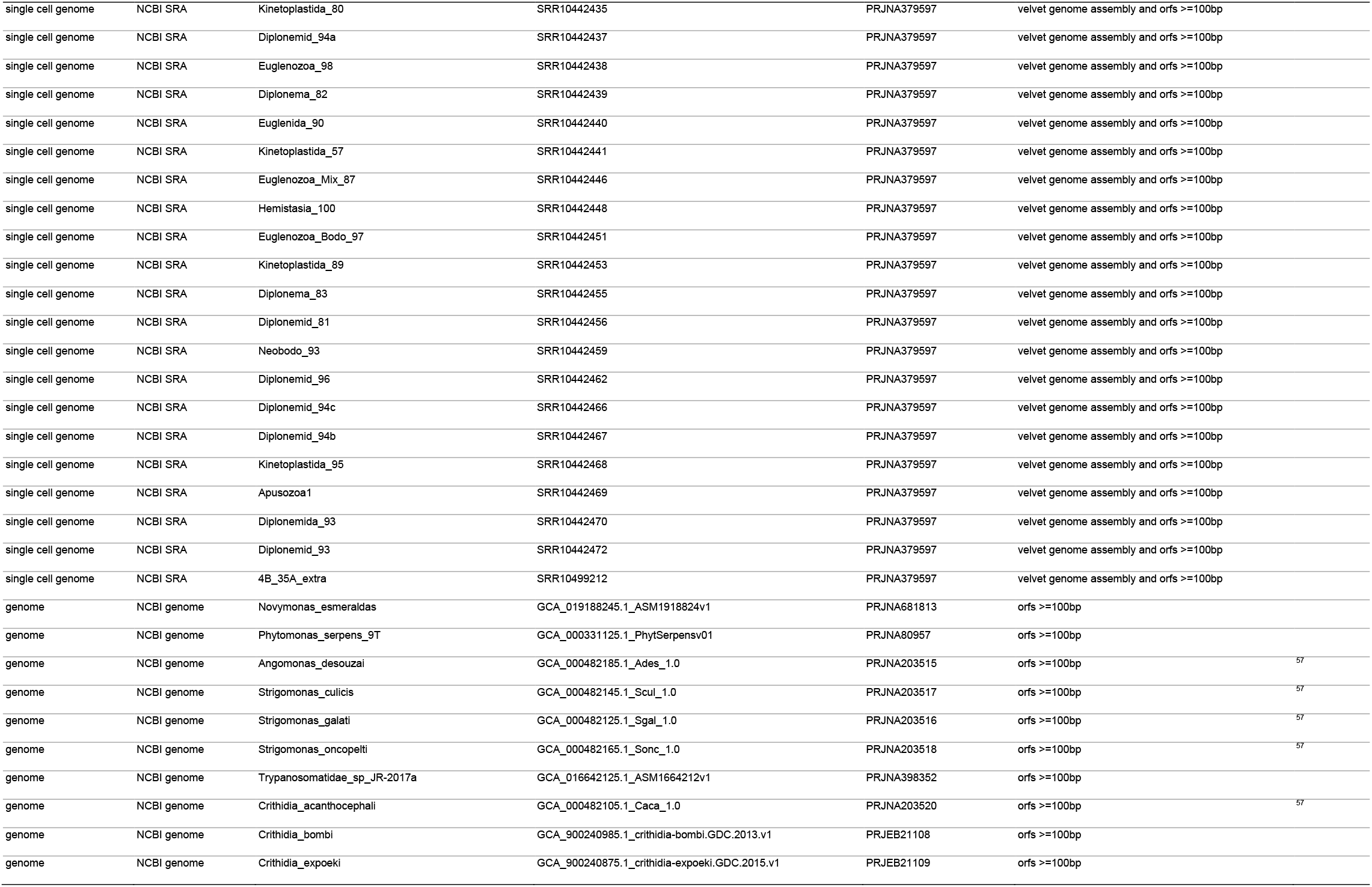
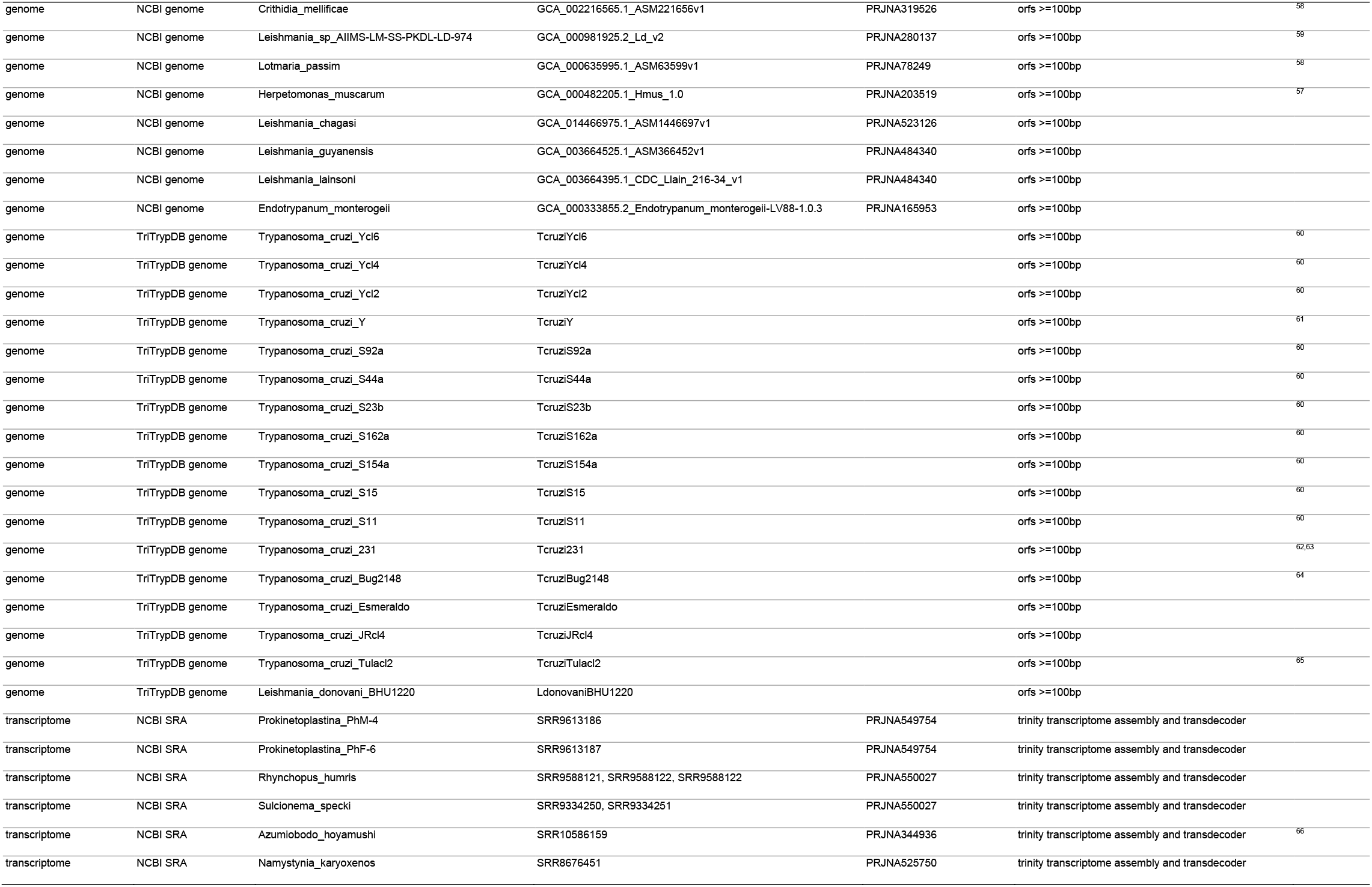
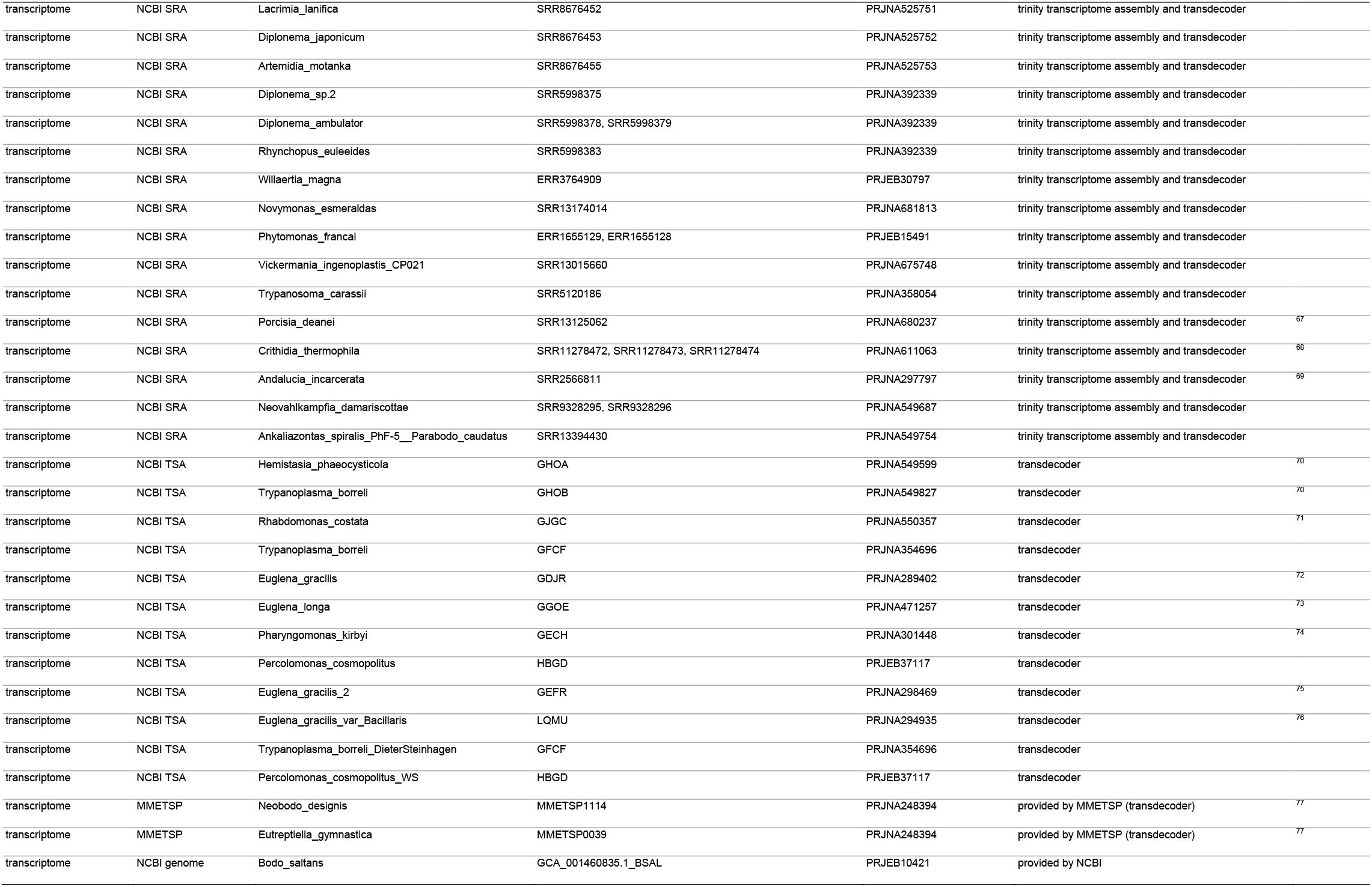
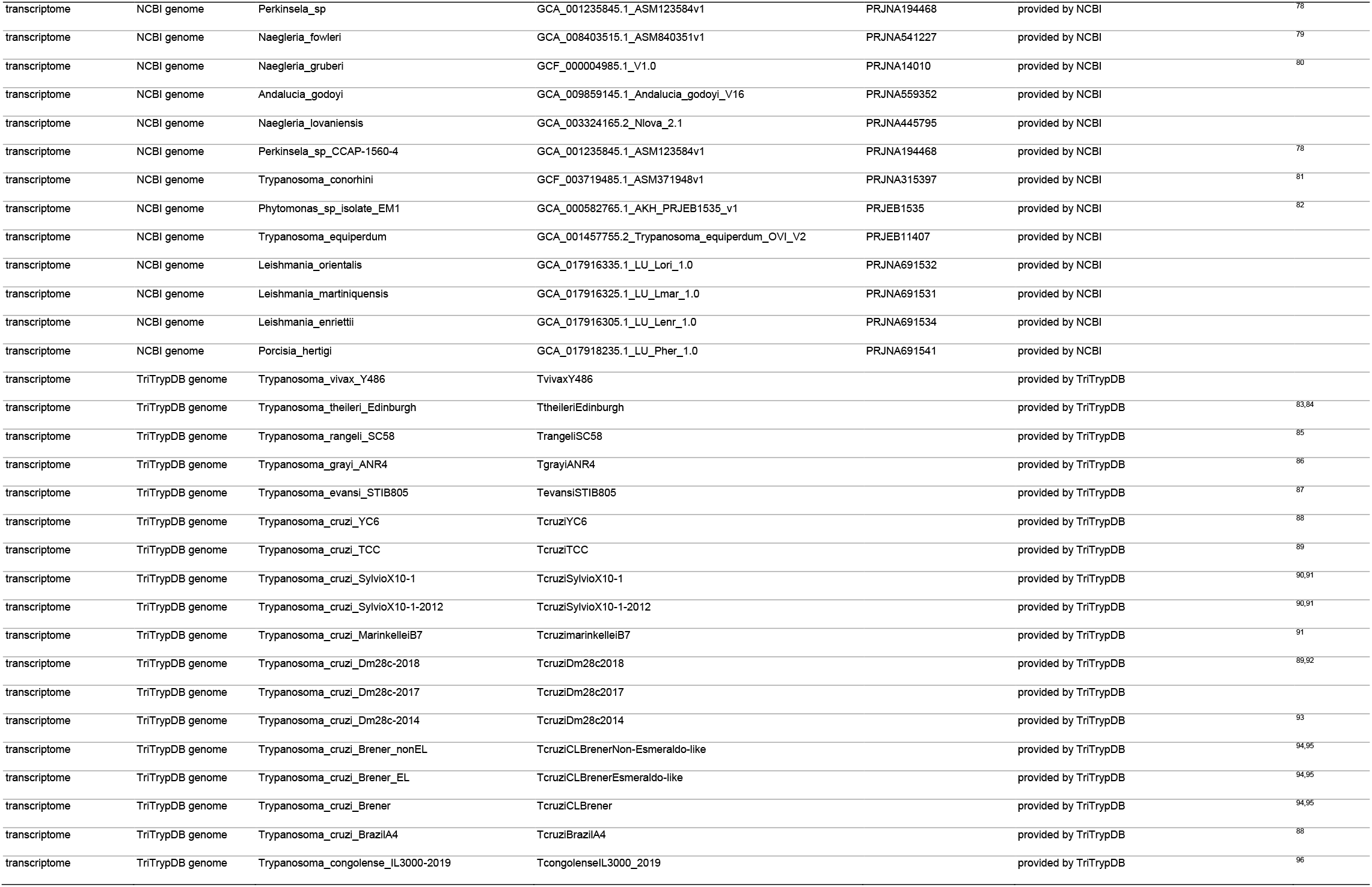
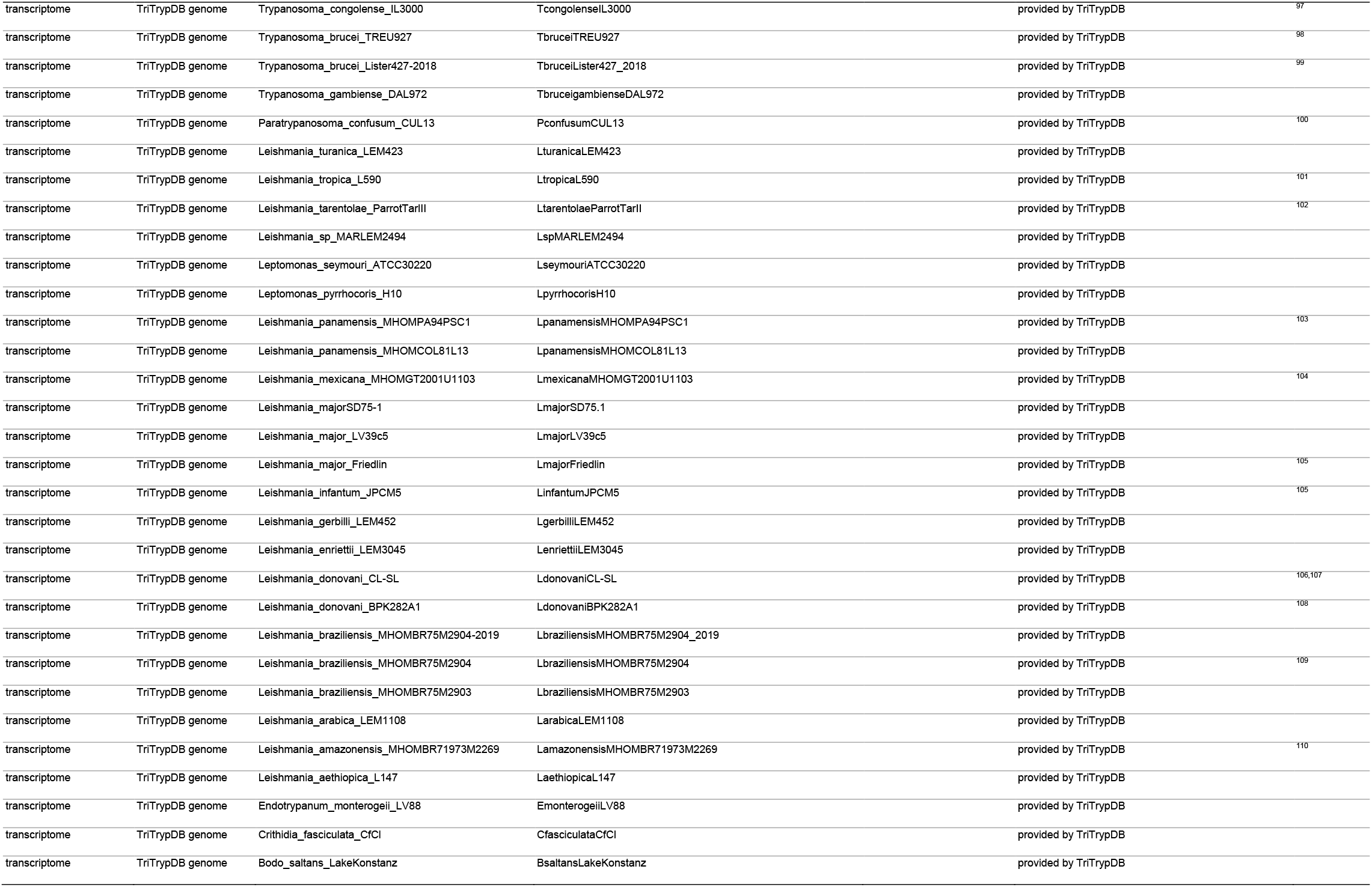
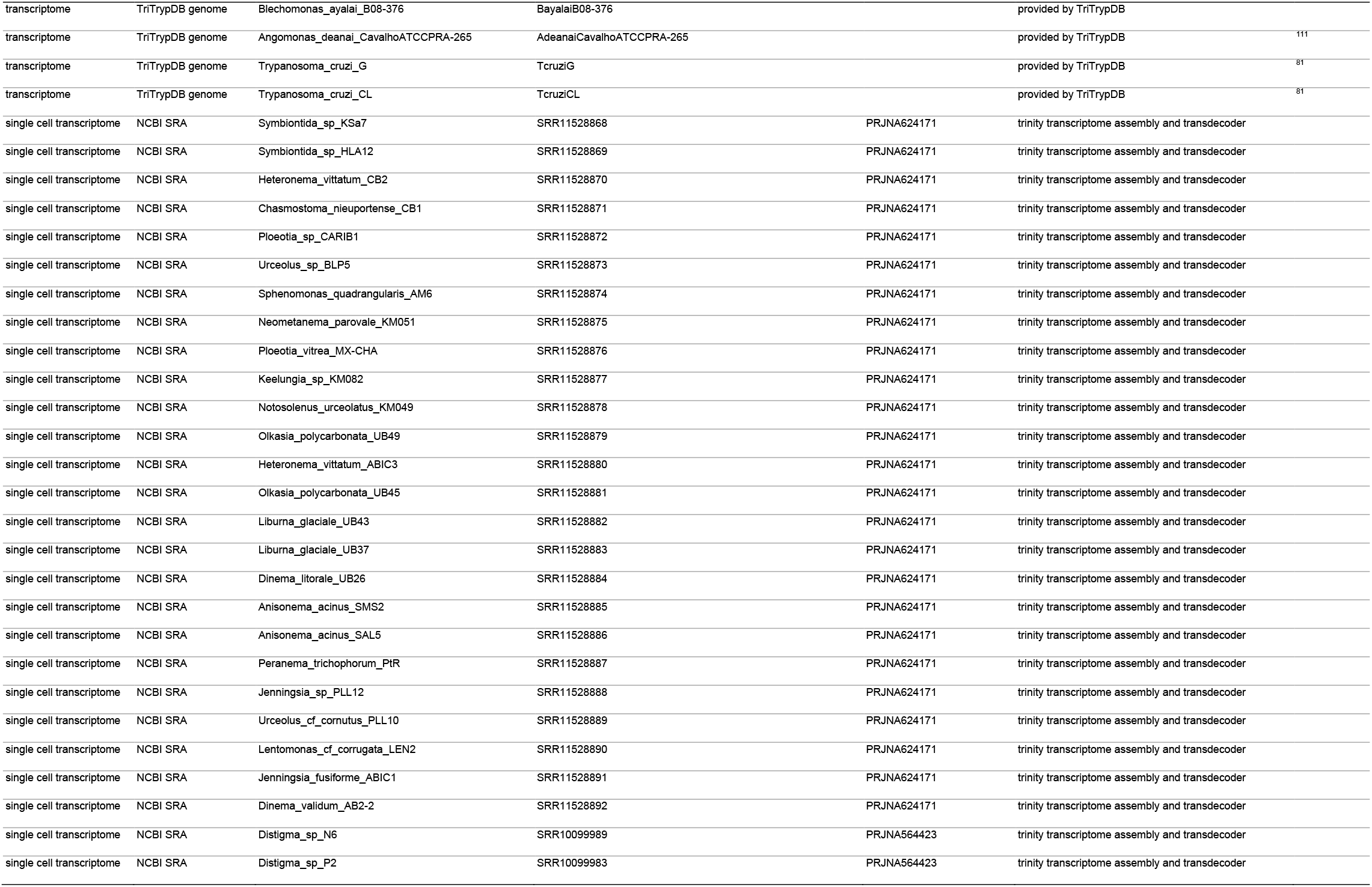
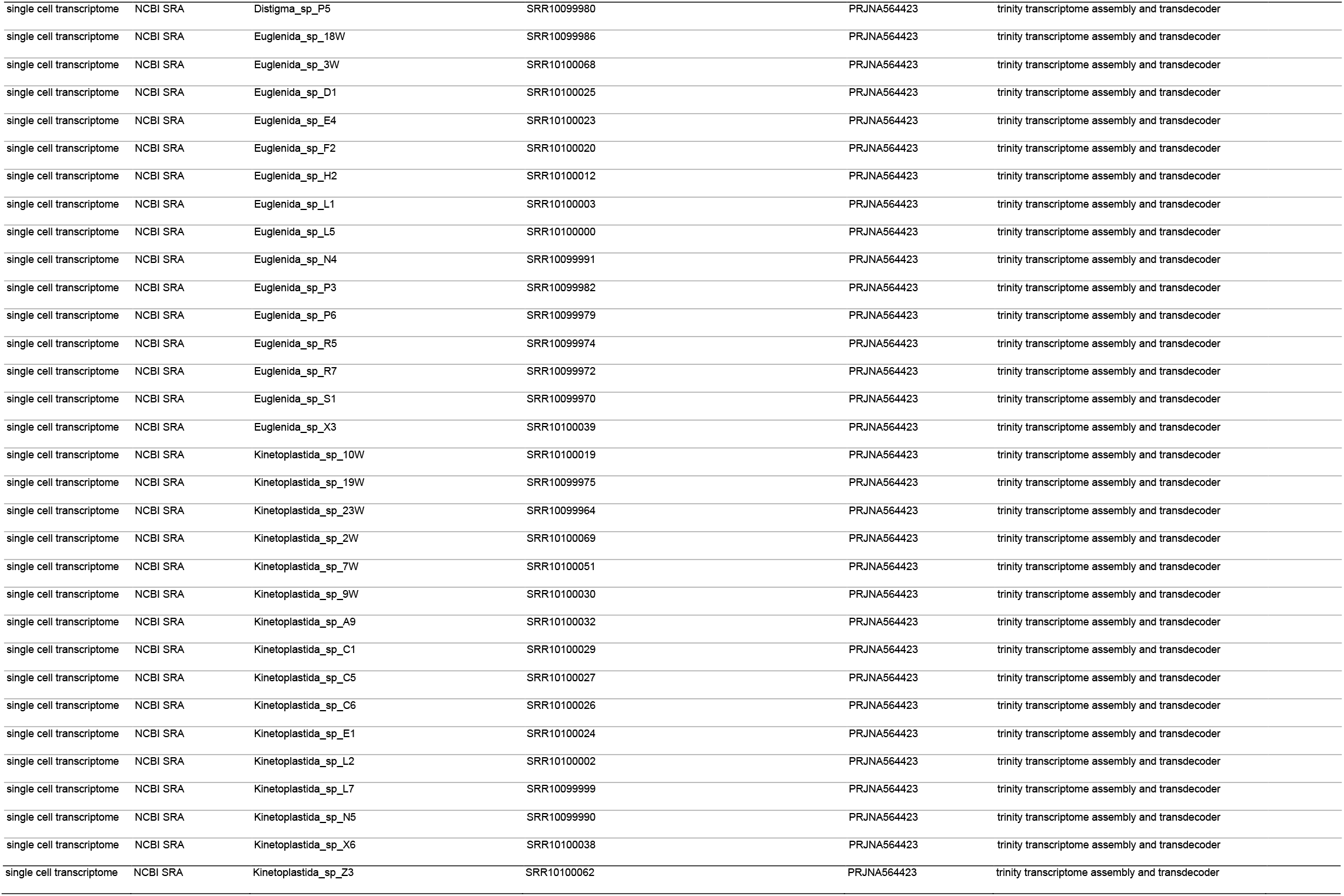
Genome and transcriptome data used to generate the Discoba protein database.

